# Characterization of a fungal mixed-linkage glucan synthase

**DOI:** 10.64898/2026.09.17.752320

**Authors:** Benjamin Menn, Klaus Pfeffer, Vicente Ramírez, Markus Pauly

## Abstract

Mixed-linkage (1,3;1,4)-β-glucan (MLG) is a cell wall polysaccharide found in fungi, bacteria, and plants, yet the enzymatic machinery and structural determinants governing fungal MLG biosynthesis remain poorly understood. In *Aspergillus fumigatus*, TFT1 (*Af*TFT1) has previously been implicated in MLG biosynthesis, but its direct biochemical activity remained unresolved. The structural characterization of *A. fumigatus* wall MLG revealed a distinct polymer profile. Heterologous expression of *Af*TFT1 in *Komagataella phaffii* enabled MLG production, generating an oligosaccharide profile that closely matched that of the native fungal polymer and demonstrating that *Af*TFT1 functions as an MLG synthase. Phylogenetic analysis placed *Af*TFT1 within a distinct glycosyltransferase family 2 (GT2) fungal lineage containing related candidate MLG synthases across diverse filamentous Ascomycota and separate from the major plant and bacterial synthase lineages. Structure-guided mutagenesis showed that individual substitutions within the transmembrane pore and switch motif altered the lichenase-derived oligosaccharide profile while retaining detectable MLG production, whereas replacement of the entire switch motif with the corresponding *Hv*CSLF6 sequence resulted in no detectable MLG production. Together, these findings establish *Af*TFT1 as a fungal MLG synthase and identify the switch motif and adjacent transmembrane region as important determinants of MLG synthase function and product fine structure, providing insight into the structural and evolutionary diversification of MLG biosynthesis.

## Introduction

Mixed-linkage glucan (MLG), also referred to as (1,3;1,4)-β-glucan, is a non-cellulosic β-glucan polymer composed of β-(1,4)-linked glucosyl residues interrupted by single β-(1,3)-linkages. MLG has been identified across diverse taxonomic groups, including bacteria (Pérez-Mendoza et al. 2015; Lampugnani et al. 2024), algae (Eder et al. 2008; Lechat et al. 2000), oomycete (Rebaque et al. 2021), fungi (Fontaine et al. 2000; Honegger and Haisch 2001; Pettolino et al. 2009) and plants (Woodward et al. 1983; Sørensen et al. 2008; Burton and Fincher 2009; Harholt et al. 2012). Despite its widespread occurrence, the molecular mechanisms underlying MLG biosynthesis and the evolutionary relationships among its synthesizing enzymes, MLG synthases, remain incompletely understood.

The presence, abundance, and structure of MLG can be assessed using complementary approaches. Immunolabelling with antibodies specific towards (1,3;1,4)-β-glucan provides qualitative information on the presence and spatial distribution of MLG in biological material (Meikle et al. 1994), whereas enzymatic hydrolysis with the enzyme lichenase enables quantitative analysis and characterization of the polymer fine structure. Lichenase specifically cleaves β-(1,4)-linkages adjacent to β-(1,3)-linked glucosyl residues, generating oligosaccharides that reflect the arrangement of linkages within the MLG polymer (McCleary 1988). Depending on the polymer structure, lichenase digestion generates oligosaccharides of different degrees of polymerization (DP), including DP2 (G3G), DP3 (G4G3G), DP4 (G4G4G3G), and DP5 (G4G4G4G3G), where G denotes glucose and the numbers represent the glycosidic linkages. The relative molar abundance of these products provides a characteristic fingerprint of MLG fine structure and can therefore be used to compare polymers produced by different synthases or organisms (Chang et al. 2021).

MLG biosynthesis has been characterized in greater detail in plants and bacteria. In plants, MLG synthase activity has been demonstrated for members of the cellulose synthase-like (CSL) family of glycosyltransferase family 2 (GT2) proteins, including members from the *CSLF, CSLH*, and *CSLJ* subfamilies (Burton et al. 2006; Doblin et al. 2009; Little et al. 2018; Purushotham et al. 2022). Plant MLG exhibits variation in fine structure, which is commonly characterized by the relative abundance of lichenase-derived DP3 and DP4 oligosaccharides, with the DP3:DP4 molar ratio used to describe differences among polymers (Burton and Fincher 2009; Jobling 2015; Chang et al. 2021). MLG synthases have also been identified and characterized in bacteria (Chang et al. 2023; Lampugnani et al. 2024; Saldivar et al. 2026), where MLG structures include polymers that generate predominantly DP2 (Pérez-Mendoza et al. 2015) or distinct combinations of products such as DP3 and DP5 (Lampugnani et al. 2024). Sequence-level determinants underlying these MLG structural variations have been investigated primarily in plants. In barley, biochemical and structural studies of the MLG synthase *Hv*CSLF6 have identified individual amino acid residues and a conserved switch motif that influence synthase activity and can modulate the fine structure of the resulting polymer (Jobling 2015; Dimitroff et al. 2016; Purushotham et al. 2022).

In contrast, although MLG has been detected in several fungal species, its biosynthesis remains comparatively poorly understood. In *Aspergillus fumigatus* (*A. fumigatus*), MLG has been detected in its wall (Fontaine et al. 2000), and genetic deletion of a putative MLG synthase, *A. fumigatus* TFT1 (*Af*TFT1), was associated with loss of detectable MLG (Samar et al. 2015). These findings implicated *Af*TFT1 in MLG biosynthesis, but did not establish whether the protein directly functions as an MLG synthase or what the fine structure of the produced MLG is. Moreover, the evolutionary relationship of *Af*TFT1 to characterized MLG synthases and the distribution of related proteins across fungi remain unclear.

Here, we investigated the biochemical function and structural determinants of *A. fumigatus* TFT1. By combining heterologous expression in *Komagataella phaffii* (formerly *Pichia pastoris*), comparative phylogenetics, and structure-guided mutagenesis based on structural alignment with plant *Hv*CSLF6, we establish *Af*TFT1 as a functional fungal MLG synthase, define its characteristic MLG fine structure, and identify sequence features within structurally conserved regions that influence product composition.

## Materials and Methods

### *Aspergillus fumigatus* growth conditions

*A. fumigatus* was maintained on Sabouraud-Dextrose-Agar (Mast Diagnostica GmbH, Reinfeld, Germany). For liquid cultures, conidia were inoculated into Sabouraud dextrose broth (4% dextrose and 1% peptone, pH 5.6) and incubated at 37 °C with shaking at 220 rpm for 24 h, following previously described growth conditions (https://www.aspergillus.org.uk/lab_protocols/growth-and-storage-of-aspergillus-fumigatus/). The resulting hyphal biomass was harvested by centrifugation, the culture medium was removed, and the biomass was resuspended in 100% ethanol.

### Plasmid construction, selection, and generation of *Af*TFT1 variants

The *Af*TFT1 coding sequence (NCBI accession XP_748682.1) was synthesized with flanking *Bsp*119I and *Xho*I restriction sites (GeneArt Thermo Fisher Scientific) and cloned into the *P. pastoris* (*Komagataella phaffii*) expression vector pPICZB (Invitrogen) by restriction-ligation cloning. The synthesized *Af*TFT1 fragment was amplified using the *Af*TFT1_F1/*Af*TFT1_R1 primer pair (Supplementary Table S1). The pPICZB vector and the *Af*TFT1 amplicon were subsequently digested with Bsp119I and XhoI (Thermo Fisher Scientific), purified, and ligated using T4 DNA ligase (New England Biolabs). The resulting plasmid was transformed into *E. coli* TOP10F’ cells, propagated, and verified by whole-plasmid sequencing.

The *Af*TFT1 protein structure was predicted using ColabFold (Mirdita et al. 2022) through the AlphaFold prediction tool implemented in UCSF ChimeraX (Meng et al. 2023) and structurally aligned with the experimentally determined structure of *Hv*CSLF6 (Purushotham et al. 2022) using the Matchmaker tool in UCSF ChimeraX. Residues occupying equivalent structural positions were identified based on three-dimensional structural overlap. Based on this comparison, a switch motif replacement variant (*Af*TFT1^SM*Hv*CSLF6^) and four single-amino-acid substitution variants (*Af*TFT1^C524I^, *Af*TFT1^I519S^, *Af*TFT1^I551Y^, and *Af*TFT1^G392D^) were generated by replacing selected *Af*TFT1 residues with the corresponding residues from *Hv*CSLF6.

For generation of the switch motif replacement variant *Af*TFT1^SM*Hv*CSLF6^, a synthetic *Af*TFT1 fragment was ordered in which the designated native switch motif sequence (residues 548-558, APYIALCIVRS) was replaced by the corresponding *Hv*CSLF6-derived sequence (residues 781-791, TASCSAYLAAV).

The synthetic fragment was amplified by PCR using the *Af*TFT1^SM*Hv*CSLF6^_F1/*Af*TFT1^SM*Hv*CSLF6^_R1 primer pair (Supplementary Table S1). The native switch motif region from *Af*TFT1 was excised from the pPICZB+*Af*TFT1 plasmid by digestion with KpnI and LguI (Thermo Fisher Scientific), and *Af*TFT1^SM*Hv*CSLF6^ insert was digested accordingly. Subsequently, vector backbone and insert were purified and ligated using T4 DNA ligase (New England Biolabs). The resulting plasmid was transformed into *E. coli* TOP10F’ cells, propagated, and verified by whole-plasmid sequencing.

The single amino acid substitution variants (*Af*TFT1^C524I^, *Af*TFT1^I519S^, *Af*TFT1^I551Y^, and *Af*TFT1^G392D^) were generated using a two-fragment PCR-based site-directed mutagenesis strategy. For each variant, two overlapping amplicons spanning the entire plasmid were generated using the constant vector backbone primers Variant_Bb_Fw and Variant_Bb_Rv in combination with variant-specific mutagenic primers. The two PCRs were performed using the primer pairs Variant_Bb_Fw/VariantX_Rv and VariantX_Fw/Variant_Bb_Rv, where X denotes the respective substitution. As an example, the *Af*TFT1^C524I^ variant was generated using the primer combinations Variant_Bb_Fw/VariantC524I_Rv and VariantC524I_Fw/Variant_Bb_Rv (Supplementary Table S1). The variant-specific primers introduced the desired nucleotide mismatches and generated complementary overlapping sequences carrying the mutations. The resulting overlapping fragments were assembled into the full-length mutant plasmid using Gibson Assembly Master Mix (New England Biolabs). The assembled plasmids were transformed into *E. coli* NEB® 5-alpha cells, propagated, and verified by whole-plasmid sequencing.

### *P. pastoris* (*Komagataella phaffii*) transformation and growth conditions

For yeast transformation, all generated constructs were linearized with MssI (Thermo Fisher Scientific) and integrated into the genome of *K. phaffii* strain X-33 via homologous recombination at the *AOX1* locus, following the protocol provided with the EasySelect™ *Pichia* Expression Kit (Invitrogen). Transformants were selected on YPDS agar containing Zeocin [100 µg ml^-1^]. Correct genomic integration at the *AOX1* locus was confirmed by PCR and Sanger sequencing of amplicons generated from gDNA of selected transformants using primer combinations pAOX1_F/*Af*TFT1_R2 and *Af*TFT1_F2/tAOX1_R (Supplementary Table S1).

For recombinant protein production, *K. phaffii* transformants were cultivated according to the EasySelect™ *Pichia* Expression Kit protocol (Invitrogen) with minor modifications. Cells were initially cultivated in buffered glycerol complex medium (BMGY) at 30 °C and 225 rpm for 24 h. Cells were harvested by centrifugation at 3,000 x *g* for 5 min, washed once with distilled water, and resuspended in buffered methanol-complex medium (BMMY) to induce expression. Cultures were incubated at 30 °C and 225 rpm for 72 h, with BMMY medium replaced every 24 h to maintain methanol induction.

### Barley growth conditions

Root material was collected from 7-day-old wild-type barley seedlings grown as described by (Guo et al. 2025) and provided by Dr. Li Guo (University of Bonn).

### Immunolabelling of *A. fumigatus* and *K. phaffii* cells

*A. fumigatus* hyphae and induced *K. phaffii* cultures (OD_600_ = 2) were collected by centrifugation at 3,000 x *g* for 5 min at 4 °C and washed three times with phosphate-buffered saline (PBS, pH 7.4). Cells were blocked in PBS containing bovine serum albumin (PBS-BSA; 10 mg mL^-1^) for 30 min at 4 °C with vertical rotation at 9 rpm.

Following blocking, cells were collected by centrifugation at 3,000 × *g* for 5 min at 4 °C and incubated with anti-(1,3:1,4)-β-glucan mouse IgG antibody (Meikle et al. 1994) in PBS-BSA for 1 h at 4 °C with vertical rotation at 9 rpm. Cells were washed three times with PBS and then incubated with goat anti-mouse IgG/IgM (H+L) secondary antibody conjugated to Alexa Fluor™ 488 (Jackson ImmunoResearch #115-545-044; 1:200 dilution) in PBS-BSA for 30 min at 4 °C with vertical rotation in the dark. Cells were subsequently washed three times with PBS and resuspended in PBS for fluorescence microscopy.

Fluorescence images were acquired using a Leica DM200 microscope equipped with pE-300 white illumination and an L5 filter cube (Leica Microsystems). Alexa Fluor™ 488 was detected using an excitation wavelength of 488 nm and an emission range of 500 - 530 nm.

### Preparation of alcohol-insoluble residues (AIR)

*A. fumigatus* and *K. phaffii* cells were harvested by centrifugation at 12,000 x *g* for 5 min and the pellets were immediately frozen in liquid nitrogen. Barley root material was freeze-dried. Alcohol-insoluble residue (AIR) was subsequently prepared from all samples as described by (Karbach et al. 2026)). Following preparation and drying, the resulting AIR was resuspended to a slurry in deionized water to a final concentration of 20 mg mL^-1^.

### Determination of (1,3;1,4)-β-glucan content and fine structure

The (1,3;1,4)-β-glucan content of *K. phaffii* AIR was determined using the β-glucan Assay Kit (Mixed-Linkage; K-BGLU; Megazyme) according to the manufacturer’s instructions. Briefly, 2 mg of AIR was incubated with a lichenase to hydrolyse mixed-linkage β-glucans into oligosaccharides. The resulting hydrolysate was subsequently treated with β-glucosidase to release glucose. The liberated glucose was quantified colorimetrically using a glucose oxidase/peroxidase (GOPOD) assay at 510 nm, and (1,3;1,4)-β-glucan content was calculated from the determined glucose concentration according to the manufacturer’s instructions.

For structural characterization of the synthesized (1,3;1,4)-β-glucans, 2 mg of AIR suspensions were incubated with 2 U of lichenase (endo-1,3:1,4-β-D-glucanase from *Bacillus subtilis;* Megazyme) in 50 mM MES buffer (pH 6.5) at 50 °C for 90 min. Following enzymatic hydrolysis, samples were centrifuged at 12,000 x *g* for 5 min, and the supernatants containing the released oligosaccharides were collected. Oligosaccharides were purified using Supelclean^™^ ENVI-Carb™ SPE columns (Sigma-Aldrich) according to the manufacturer’s instructions. Lichenase-released oligosaccharides were analysed by high-performance anion-exchange chromatography with pulsed amperometric detection (HPAEC-PAD) using an IC Amperometric Detector (Metrohm), as previously described (Schultink et al. 2013; Immelmann et al. 2023).

For comparison of oligosaccharide product profiles, the quantities of individual lichenase-released oligosaccharides were converted from mass to molar amounts using their respective molecular masses, and molar ratios were calculated accordingly.

### Glycosyl Linkage Analysis

Glycosyl linkage analysis was performed on oligosaccharide fractions collected after HPAEC-PAD separation. The samples were derivatized to their partially methylated alditol acetates (PMAAs). Methylation, hydrolysis, reduction, and acetylation derivatization were performed as previously described (Schultink et al. 2013; Voiniciuc et al. 2019). Separation and detection of PMAAs were performed using a 7890B GC system connected to a 5977A series Quadrupole MS system (both Agilent). GC separation was performed using a SP-2380 Fused Silica Capillary Column (Supelco). Carbohydrate peaks were annotated according to individual retention times and ion fragmentation spectra in comparison to the CCRC spectral database for PMAAs (https://glygen.ccrc.uga.edu/ccrc/specdb/ms/pmaa/pframe.html).

### Phylogenetic analysis of MLG synthases

To determine the evolutionary relationship of *Af*TFT1 to previously characterized MLG synthases, *Af*TFT1 was compared with representative MLG synthases selected based on published functional characterization (Supplementary Table S2). Protein sequences were aligned using MAFFTv7 with the L-INS-i method (Katoh et al. 2005) and trimmed using trimAl (Capella-Gutiérrez et al. 2009). The best-fitting amino acid substitution model was determined using ModelFinder (Kalyaanamoorthy et al. 2017). Maximum-likelihood phylogenetic inference was performed using IQ-TREE 2 (Minh et al. 2020), and branch support was assessed using 1,000 ultrafast bootstrap replicates (Hoang et al. 2018). The resulting phylogenetic tree was visualized and annotated using iTOL v6 (Letunic and Bork 2024).

### Phylogenetic analysis of *Af*TFT1-related GT2 proteins

To investigate the distribution of *Af*TFT1-related GT2 proteins and identify candidate proteins for future functional characterization, the *Af*TFT1 protein sequence was used as query in a BLASTP search. Hits with bit scores > 160 were selected for further analysis, and manually curated by removing exact amino-acid duplicates and incomplete sequences based on a multiple-sequence alignment. Functionally characterized GT2 proteins representing fungal, plant, and bacterial glucan synthases were included as reference sequences (Supplementary Table S3). Maximum-likelihood phylogenetic inference was performed using IQ-TREE 2 (Minh et al. 2020), and branch support was assessed using 5,000 ultrafast bootstrap replicates. The tree was visualized and annotated using iTOL v6 (Letunic and Bork 2024).

## Results

### Characterization of mixed-linkage β-glucan in *Aspergillus fumigatus* reveals a distinct polymer fine structure

The presence and fine structure of mixed-linkage β-glucan (MLG) in *A. fumigatus* were characterized using complementary immunolabelling and enzymatic approaches. For this purpose, fungal hyphae were immunolabelled using a monoclonal anti-MLG antibody. Strong fluorescence was observed predominantly at the hyphal surface (Fig. 1A), indicating the presence of antibody-accessible MLG within the fungal wall. These results are consistent with previous observations in *A. fumigatus* (Samar et al. 2015).

**Figure 1.**
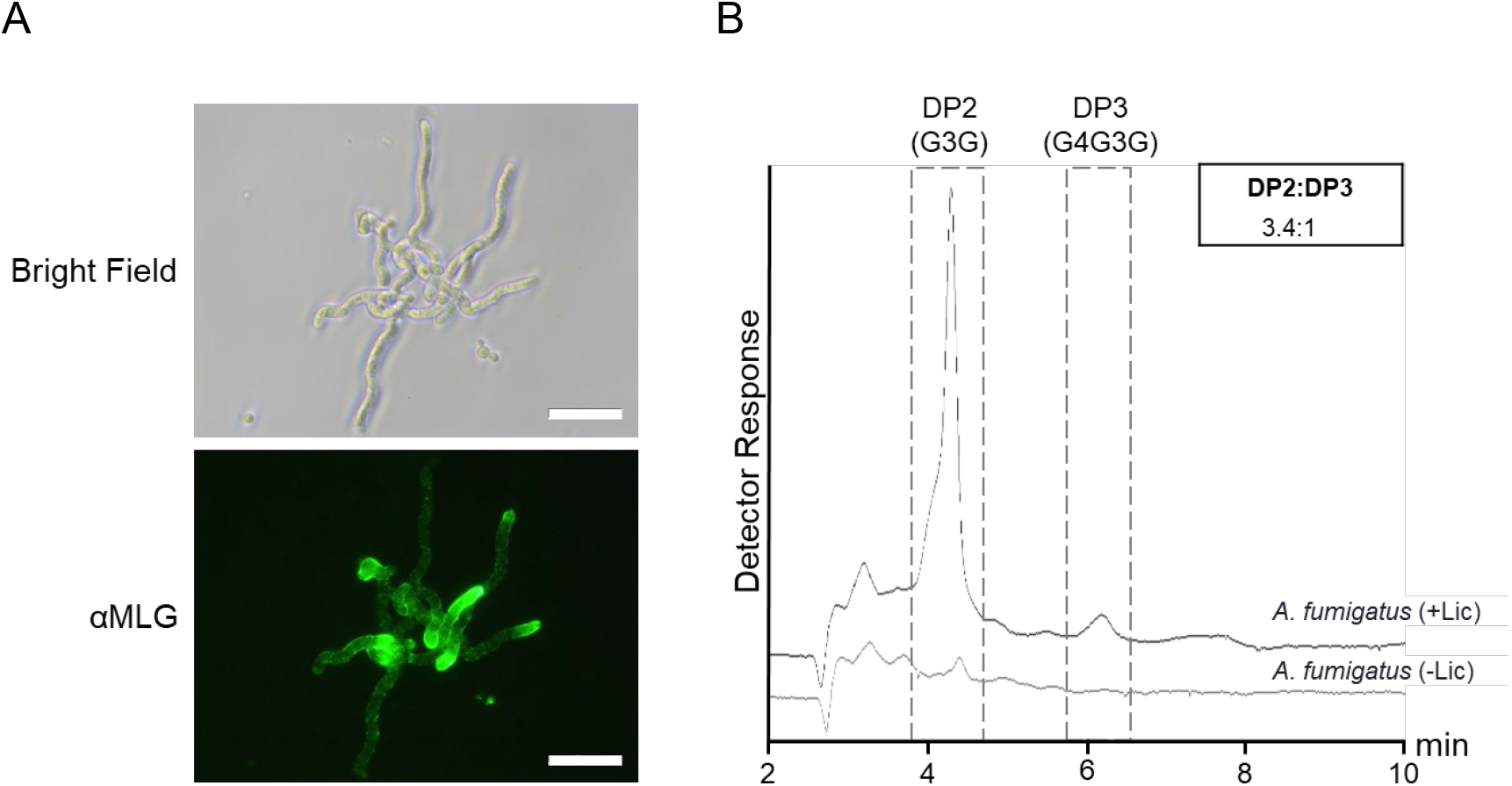
Characterization of mixed-linkage glucan (MLG) in *Aspergillus fumigatus*. (A) Representative microscopy images of *A. fumigatus* hyphae following immunolabelling using an anti-MLG (aMLG) monoclonal antibody. Scale bars = 50 µm. (B) Representative HPAEC-PAD chromatograms of lichenase-treated (*A. fumigatus* +Lic) and untreated (*A. fumigatus* −Lic) AIR preparations. Released oligosaccharides were identified by comparison with the retention times of MLG-oligosaccharide standards and quantified relative to an internal standard. Inset value indicates the DP2:DP3 molar ratio of the detected oligosaccharides. DP, degree of polymerization.

For structural characterization of the MLG polymer, wall material (alcohol-insoluble residue; AIR) isolated from *A. fumigatus* was hydrolysed with lichenase, and the released oligosaccharides were analysed by HPAEC-PAD (Fig. 1B). Lichenase-treated *A. fumigatus* AIR (+Lic) generated distinct peaks co-eluting with MLG standards DP2 (G3G) and DP3 (G4G3G). These oligosaccharides were absent from non-digested control samples (-Lic). Quantification of chromatographic peak areas revealed relative abundances of 0.11 ± 0.01 µg DP2 and 0.04 ± 0.01 µg DP3 per mg of lichenase-hydrolysed AIR, corresponding to a DP2:DP3 molar ratio of 3.4:1. Together, these data provide evidence for the presence of a structurally defined MLG in *A. fumigatus* walls, prompting further investigation of the enzyme responsible for its biosynthesis, *Af*TFT1.

### Phylogenetic analysis places *Af*TFT1 in a previously uncharacterized fungal GT2 lineage

Two complementary phylogenetic analyses were performed to assess the relationship of *Af*TFT1 to characterized MLG synthases and to identify additional *Af*TFT1-related GT2 proteins. The first analysis focused on functionally characterized MLG synthases to determine the phylogenetic position of *Af*TFT1 relative to experimentally confirmed enzymes. The resulting phylogenetic tree revealed distinct clades that largely correspond to the taxonomic origin of the analysed MLG synthases (Fig. 2). Plant and bacterial MLG synthases formed separate clades, whereas *Af*TFT1 occupied a distinct lineage among the characterized MLG synthases.

**Figure 2.**
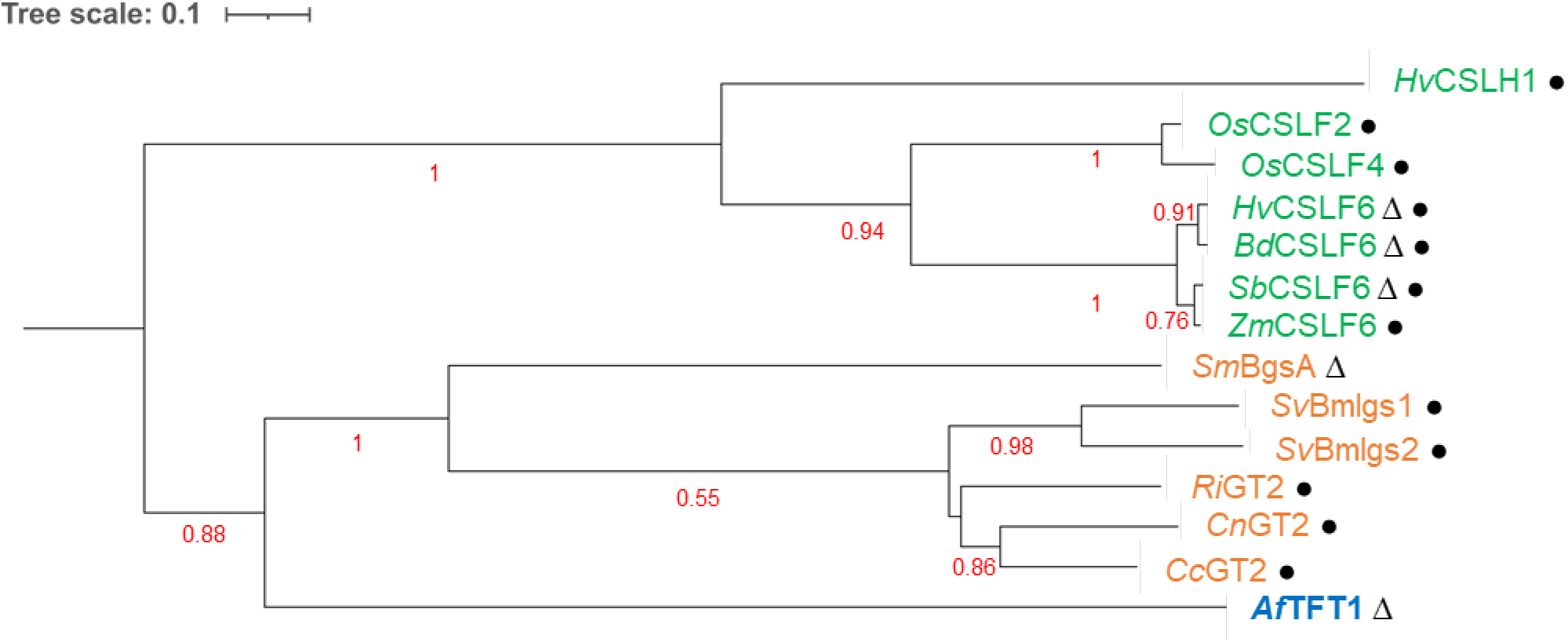
Phylogenetic relationship of *Af*TFT1 and other characterized MLG synthases. Maximum-likelihood phylogenetic tree inferred from amino acid sequences of representative plant (green), bacterial (orange) and fungal (blue) MLG synthases. The scale bar indicates the expected number of amino acid substitutions per site. Numbers at the nodes indicate bootstrap support values expressed as proportion of 1,000 pseudoreplicates. Symbols indicate experimental evidence associated with the corresponding MLG synthase protein: Δ - genetic evidence from a mutant line or strain lacking detectable MLG; • - biochemical characterization through heterologous expression. Protein accession numbers (NCBI/PDB IDs) are provided in Supplementary Table S2. Af, *Aspergillus fumigatus*; Bd, *Brachypodium distachyon*; Cc, *Clostridium cuniculi*; Cn, *Clostridium nigeriense*; Hv, *Hordeum vulgare*; Os, *Oryza sativa*; Ri, *Romboutsia ilealis*; Sb, *Sorghum bicolor*; Sm, *Sinorhizobium meliloti*; Sv, *Sarcina ventriculi*; Zm, *Zea mays*.

A second phylogenetic analysis examined a broader set of GT2 proteins identified by sequence similarity to *Af*TFT1 (Supplementary Fig. S1) to investigate the evolutionary distribution of *Af*TFT1-related proteins. The analysis revealed a distinct *Af*TFT1-like fungal lineage spanning diverse filamentous Ascomycota, including Eurotiomycetes, Sordariomycetes, Leotiomycetes, Dothideomycetes, and Pucciniomycetes, clearly separated from the major plant, bacterial, and oomycete GT2 lineages. The proteins within this *Af*TFT1-like lineage shared 70–96% amino-acid sequence identity and 80–97% sequence similarity with *Af*TFT1, further supporting their close evolutionary relationship. Their occurrence across multiple fungal classes indicates that the *Af*TFT1-like lineage extends beyond *Aspergillus* and represents a broader fungal GT2 lineage. Although MLG synthase activity has not yet been experimentally established for these proteins, their close relationship to *Af*TFT1 makes them candidates for additional fungal MLG synthases. Notably, MLG has been reported in *Pyrenophora teres* (Backes et al. 2020; Chang et al. 2021), for which the responsible biosynthetic enzyme remains unknown. Thus, the *P. teres* GT2 protein included in our analysis (entry 66), together with the *Af*TFT1-like proteins identified here, provide a set of candidates for experimental investigation of fungal MLG synthase activity.

### Heterologous expression of *Af*TFT1 enables mixed-linkage β-glucan production

*Af*TFT1 has previously been implicated in MLG biosynthesis, as *AfTFT1*-mutant *A. fumigatus* strains lacked MLG based on anti-MLG antibody labelling (Samar et al. 2015). To determine whether *Af*TFT1 can produce MLG in a heterologous cellular context, *Af*TFT1 was recombinantly expressed in *K. phaffii* (formerly *Pichia pastoris*), a host lacking detectable endogenous MLG (Fig. 3). An *Af*TFT1 expression cassette was integrated into the *K. phaffii* genome under control of the methanol-inducible *AOX1* promoter. Following induction of recombinant protein expression, *K. phaffii* cells were immunolabelled using an anti-MLG antibody. While anti-MLG antibody signal remained absent in wild-type *K. phaffii* cells, *Af*TFT1-expressing cells showed strong fluorescence at the cell surface (Fig. 3A), demonstrating the production of a polysaccharide recognized by the anti-MLG antibody.

**Figure 3.**
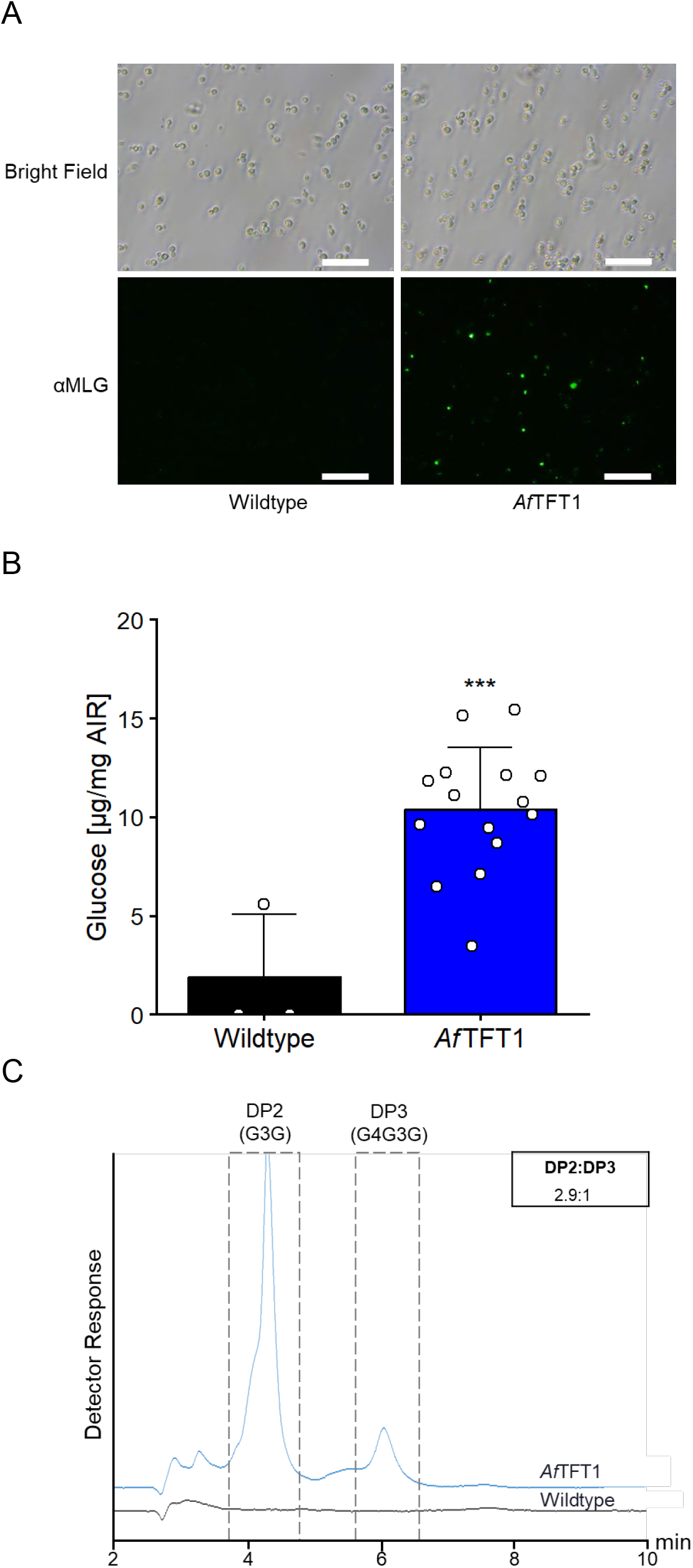
Biochemical characterization of *Af*TFT1 in *K. phaffii*. (A)Representative microscopy images of *Af*TFT1-expressing *K. phaffii* cells following immunolabelling using an anti-MLG monoclonal antibody (aMLG). Wild-type *K. phaffii* is shown as a control. Scale bars = 50 µm.(B) Detection of β-glucan derived glucose in AIR preparations from *K. phaffii* cells expressing *Af*TFT1. Data represent mean ± SD with individual technical replicates indicated by dots *(n* = 3 for wildtype and *n* = 15 for *Af*TFT1, respectively). Only positive values were considered, while negative values were set to 0. *S*tatistical significance was assessed using an unpaired Student’s t-test (*** = *P* < 0.001). C) Representative HPAEC-PAD chromatograms of lichenase-hydrolysed AIR preparations of *Af*TFT1-expressing *K. phaffii* cells. AIR from wild-type *K. phaffii* was included as a control. Released oligosaccharides were identified by comparison to the retention times of MLG-oligosaccharide standards and quantified relative to an internal standard. Inset value indicates the DP2:DP3 molar ratio of the detected oligosaccharides. DP, degree of polymerization; SD, standard deviation.

To further assess MLG production, AIR was isolated from wild-type and *Af*TFT1-expressing *K. phaffii* cells and MLG content was assayed. Wild-type *K. phaffii* AIR contained only trace levels of lichenase-released glucose (1.9 ± 3.2 µg mg^-1^ AIR; Fig. 3B), while *Af*TFT1-expressing cell AIR contained substantially higher levels (10.4 ± 3.2 µg mg^-1^ AIR), demonstrating the production of MLG.

Lichenase-released oligosaccharides were further analysed by HPAEC-PAD. No detectable oligosaccharide peaks were observed in lichenase-treated wild-type *K. phaffii* AIR, whereas analysis of lichenase-hydrolysed AIR of *Af*TFT1-expressing cells yielded two peaks that co-eluted with oligosaccharide standards for MLG DP2 (G3G) and MLG DP3 (G4G3G; Fig. 3C). Quantification of the released oligosaccharides yielded a DP2:DP3 molar ratio of 2.9:1 (Supplementary Table S5). The contents of the DP2 and DP3 peaks (see Fig. 3C) were collected as individual fractions and subjected to glycosyl linkage analysis, which confirmed identity of the detected oligosaccharide structures (Supplementary Table S4).

Taken together, these results demonstrate that *Af*TFT1 encodes a functional MLG synthase capable of directing MLG biosynthesis in a heterologous host. Expression of *Af*TFT1 in *K. phaffii* thus provides a platform for the production of MLG. In addition, functional and structure-function analyses of *Af*TFT1 and potentially other MLG synthases can be pursued using this heterologous system.

### Structure-guided mutagenesis identifies residues influencing mixed-linkage β-glucan biosynthesis and polymer fine structure in *Af*TFT1

Previous characterization of the barley MLG synthase *Hv*CSLF6 expressed in Sf9 insect cells and analysed *in vitro* showed production of an MLG polymer yielding DP3 (G4G3G) and DP4 (G4G4G3G) oligosaccharides at a molar ratio of 1:2 (Purushotham et al. 2022). To benchmark plant MLG fine structure under the analytical conditions used in this study, we analysed lichenase-released oligosaccharides of barley root AIR by HPAEC-PAD. This revealed DP2, DP3 and DP4 at a molar ratio of 1:16.2:3.4, with DP3 and DP4 as the dominant oligosaccharides (Supplementary Fig. S3; Supplementary Table S5). In contrast, both native *A. fumigatus* MLG and recombinant *Af*TFT1-produced MLG in *K. phaffii* yielded DP2 and DP3 oligosaccharides at a molar ratio of DP2:DP3 of 3.4:1 and 2.9:1, respectively. No DP4 oligosaccharides were detected under the conditions used here. Thus, *A. fumigatus* and recombinant *Af*TFT1-produced MLG exhibit a lichenase-derived oligosaccharide profile distinct from that observed for barley MLG, characterized by a high relative abundance of DP2 and the absence of detectable DP4.

Given the distinct MLG profiles produced by the fungal and plant systems, we next asked whether protein domains of specific amino acids in *Af*TFT1 corresponding to those previously implicated in MLG fine-structure control in *Hv*CSLF6 could influence the fine structure of the fungal MLG polymer. In the absence of experimentally determined fungal MLG synthase structures, the cryo-EM structure of the barley MLG synthase *Hv*CSLF6 was used as a structural reference. Previous structure-function studies of *Hv*CSLF6 identified individual amino acid residues and a conserved switch motif that influence the MLG product fine structure, including the relative abundance of lichenase-derived oligosaccharides (Jobling 2015; Purushotham et al. 2022). Structural alignment of the AlphaFold-predicted *Af*TFT1 model with the cryo-EM structure of the barley MLG synthase *Hv*CSLF6 revealed conservation of the core protein architecture despite divergence in peripheral regions (Fig. 4A). The alignment identified a structurally conserved region spanning 237 residues, with a root-mean-square-deviation (RMSD) of 1.111 Å and 29.5% sequence identity. Based on this structural correspondence, we identified a region in *Af*TFT1 corresponding to the *Hv*CSLF6 switch motif, which has previously been implicated in determining MLG fine structure, together with neighbouring residues selected for functional analysis.

**Figure 4.**
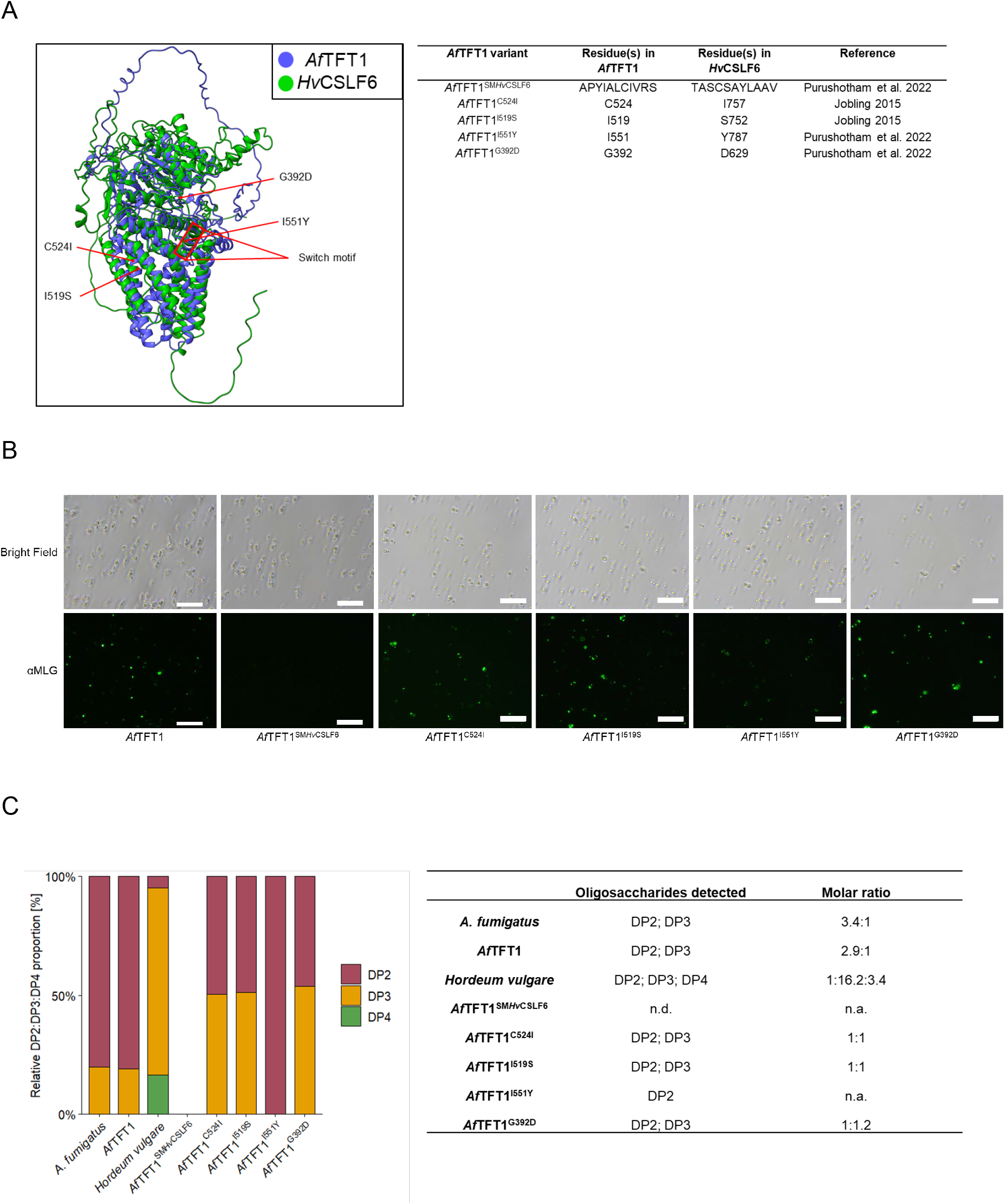
Biochemical characterization of *Af*TFT1 variants in *K. phaffii*. (A) Structure-guided *Af*TFT1 variant generation. Structural alignment of predicted *Af*TFT1 (NCBI accession XP_748682) with the cryo-EM structure of *Hv*CSLF6 (pdb ID: 8DQK) for identification of *Af*TFT1-residues corresponding to previously mutagenized positions in *Hv*CSLF6. Right side: Summary and structural details of the generated *Af*TFT1 variants. (B) Representative microscopy images of *K. phaffii* cells expressing *Af*TFT1 and individual *Af*TFT1 variants following immunolabelling using an anti-MLG monoclonal antibody. Scale bars = 50 µm. (C) Relative distribution of lichenase-released oligosaccharides from *Aspergillus fumigatus, Hordeum vulgare*, and *Komagataella phaffii* cells expressing *Af*TFT1 variants. Total released oligosaccharides were normalized to 100% for each biological sample. Coloured bars represent the relative contribution of DP2, DP3 and DP4 to the total released oligosaccharides. The five rightmost bars represent *Af*TFT1 variants, including *Af*TFT1 containing the *Hv*CSLF6 switch motif (SM*Hv*CSLF6), and the single-amino-acid substitutions (C524I, I519S, I551Y and G392D). Values in the table indicate the corresponding oligosaccharide molar ratios for each sample. DP, degree of polymerization; n.a., not applicable; n.d., not detected.

Two complementary mutagenesis strategies were employed. First, the predicted *Af*TFT1 switch motif was replaced with the corresponding motif from *Hv*CSLF6. Second, four *Af*TFT1 amino acids occupying structurally corresponding positions to *Hv*CSLF6 residues previously shown to influence the DP3:DP4 ratio were identified. Each of these residues was individually replaced with the corresponding *Hv*CSLF6 amino acid. Hence, five *Af*TFT1 variants were generated: the switch motif replacement variant *Af*TFT1^SM*Hv*CSLF6^ and four single-amino-acid substitution variants *Af*TFT1^C524I^, *Af*TFT1^I519S^, *Af*TFT1^I551Y^, and *Af*TFT1^G392D^.

The *Af*TFT1 variants were individually expressed in *K. phaffii* and analysed for MLG synthase activity by immunolabelling with the anti-MLG antibody. While expression of the *Af*TFT1 switch motif replacement variant (*Af*TFT1^SM*Hv*CSLF6^) did not result in detectable anti-MLG fluorescence, all four single-amino-acid substitution variants displayed detectable anti-MLG fluorescence (Fig. 4B).

To determine whether the substitutions also affected the fine structure of the synthesized MLG polymer, *K. phaffii* AIR was subjected to lichenase hydrolysis and the released oligosaccharides were analysed by HPAEC-PAD. No lichenase-released oligosaccharide peaks were detected for *Af*TFT1^SM*Hv*CSLF6^, consistent with the absence of detectable anti-MLG immunolabelling (Supplementary Fig. S2). In contrast, the *Af*TFT1^I551Y^ variant produced only DP2 under the applied analytical conditions, whereas the other three single-amino-acid substitution variants *Af*TFT1^C524I^, *Af*TFT1^I519S^, and *Af*TFT1^G392D^ produced both DP2 and DP3. Quantification revealed that all four variants yielded lower levels of lichenase-released MLG-derived oligosaccharides than wild-type *Af*TFT1 and exhibited mutation-dependent changes in the DP2:DP3 molar ratio (Fig. 4C; Supplementary Table S5). These results demonstrate that substitutions at the selected positions are associated with changes in MLG accumulation and in the distribution of lichenase-derived oligosaccharides, indicating that these regions of *Af*TFT1 contribute to its functional properties and influence the resulting MLG product fine structure.

## Discussion

MLG has previously been detected in the cell wall of *A. fumigatus*, and deletion of *Af*TFT1 resulted in loss of detectable MLG, implicating *Af*TFT1 in fungal MLG biosynthesis (Fontaine et al. 2000; Samar et al. 2015). Extending this work, we characterized *A. fumigatus* MLG and investigated the activity of *Af*TFT1 by heterologous expression. Lichenase hydrolysis of *A. fumigatus* AIR released DP2 and DP3 oligosaccharides, revealing a fine-structure profile distinct from the predominantly DP3 profile with a minor DP4 component reported for MLG produced by the bacterial synthase *Ri*GT2 (Chang et al. 2023) and the DP3- and DP4-rich profile characteristic of grass MLGs, including barley MLG synthesized by *Hv*CSLF6 (Jobling 2015; Purushotham et al. 2022). Importantly, expression of *Af*TFT1 in *K. phaffii* resulted in an MLG with same major lichenase-released oligosaccharides, DP2 and DP3, as MLG isolated from *A. fumigatus*, with similar molar ratios. Together, these results demonstrate that *Af*TFT1 is capable of synthesizing MLG with a fine-structure profile comparable to that of *A. fumigatus* MLG, extending the previous genetic evidence linking *Af*TFT1 to MLG biosynthesis in *A. fumigatus*.

Biochemical and cryo-electron microscopy of heterologously expressed HvCslF6 revealed that this MLG synthase functions as a monomer producing both β-(1,3)- and β-(1,4)-glucosidic linkages of MLG (Purushotham et al. 2022). The structural similarity of the fungal *Af*TFT1 suggests a similar mechanism whereby *Af*TFT1 alone may be sufficient for the production of MLG. However, there might be other endogenous *K. phaffii* proteins that may influence MLG abundance, localization, or polymer structure.

In *Hv*CSLF6, previous work demonstrated that polymer fine structure is influenced by specific structural elements of the enzyme near the catalytic pore and transmembrane region (Jobling 2015; Purushotham et al. 2022). Specifically, single-amino-acid substitutions at positions adjacent to the transmembrane pore – such as D629G, S752G and I757L – as well as modifications within its conserved switch motif (Y787H) systematically altered the molar ratio of DP3 to DP4 oligosaccharides, shifting synthesis toward a higher proportion of DP4 units. Furthermore, swapping the entire CSLF6 switch motif with that of a cellulose synthase (CESA) significantly reduced the frequency of β-(1,3)- linkages introduced into the polymer (Purushotham et al. 2022).

To test whether analogous structural features contribute to fungal MLG fine structure, we targeted positions in *Af*TFT1 corresponding to residues in *Hv*CSLF6 previously implicated in determining MLG fine structure. All four single substitutions (*Af*TFT1^C524I^, *Af*TFT1^I519S^, *Af*TFT1^I551Y^, and *Af*TFT1^G392D^) retained detectable MLG production but altered the DP2:DP3 molar ratio relative to non-substituted (wild-type) *Af*TFT1. Thus, individual residues within these structurally corresponding regions can be changed without stopping MLG production, while still altering the lichenase-derived oligosaccharide profile. The particularly pronounced effect of I551Y, which yielded only MLG oligosaccharides of DP2 under the applied analytical conditions, further highlights the sensitivity of the product profile to individual residue changes within this region. The location of these residues within the switch motif or adjacent transmembrane pore is consistent with models proposed by (Jobling 2015) and (Dimitroff et al. 2016), in which structural features of this region contribute to MLG fine structure.

A notable difference between the two systems was observed when the entire switch motif was replaced. In *Hv*CSLF6, mutation or replacement of the switch motif reduced the abundance of β-(1,3)- linkages in MLG while retaining enzyme activity (Jobling 2015; Purushotham et al. 2022). Here, introduction of the complete *Hv*CSLF6 switch motif into *Af*TFT1 (*Af*TFT1^SM*Hv*CSLF6^) resulted in no detectable MLG production under the conditions tested. This contrast suggests that the switch region is functionally constrained at the level of its overall sequence context, even though individual residues within the region can be altered while retaining detectable synthase activity. Thus, the switch motif and adjacent transmembrane pore may accommodate specific residue changes that modulate product composition, whereas replacement of the entire motif disrupts enzyme function. However, effects on protein stability, localization, catalytic activity, or processivity not assessed here may also contribute to the observed differences.

Phylogenetic analysis placed *Af*TFT1 into a distinct fungal GT2 lineage that is evolutionarily divergent from previously characterized plant and bacterial MLG synthases. *Af*TFT1-like proteins were identified across diverse filamentous Ascomycota lineages, including Eurotiomycetes, Sordariomycetes, Leotiomycetes, and Dothideomycetes, suggesting that this lineage is not restricted to *Aspergillus*. Based on their phylogenetic relationship to *Af*TFT1, these proteins represent candidate fungal MLG synthases, although their enzymatic activity remains to be experimentally validated. Conversely, the possibility that additional proteins within the broader fungal GT2 clade may also possess MLG synthase activity cannot be excluded. The deep evolutionary divergence between the fungal, plant, and bacterial MLG synthase lineages is accompanied by differences in the fine structure of the MLG polymers they produce. Together, these observations raise the possibility that MLG synthase activity arose independently in distinct GT2 lineages, although additional functional characterization of candidate enzymes will be required to test this hypothesis.

Overall, our findings establish *Af*TFT1 as a functional fungal MLG synthase belonging to an evolutionarily novel GT2 lineage and show that targeted substitutions within structurally conserved regions of the switch motif and adjacent transmembrane pore alter MLG accumulation and fine structure. By demonstrating MLG biosynthesis through heterologous expression of *Af*TFT1, this work expands the known diversity of MLG synthases and provides a framework for investigating the molecular and evolutionary basis of fungal MLG biosynthesis.

## Author contributions

VR and MP conceived and designed the project. BM performed experiments and analysed data. BM, KP, VR and MP wrote the manuscript. All authors contributed to the article and approved the submitted version.

## Acknowledgements

The authors would like to thank Claudia Wallscheid and Nicole Küpper for excellent technical assistance. This work was funded by the Cluster of Excellence on Plant Sciences (CEPLAS) by the Deutsche Forschungsgemeinschaft (DFG; German Research Foundation) under Germany’s Excellence Strategy-EXC 2048/1-Project ID: 390686111.

## Data availability statement

All data supporting the findings in this study are available within the paper and its Supporting Information

## Supporting information

**Figure S1**. Phylogenetic relationships of *Af*TFT1-related GT2 glycosyltransferases.

**Figure S2**. Biochemical characterization of lichenase-released oligosaccharides from *K. phaffii* expressing *Af*TFT1 variants.

**Figure S3**. Biochemical characterization of lichenase-released oligosaccharides from *Hordeum vulgare* root material.

**Table S1**. Oligonucleotide primers used in this study.

**Table S2**. Protein sequences and accession numbers used for phylogenetic analysis of *Af*TFT1 and characterized MLG synthases.

**Table S3**. BLAST search results and taxonomic classification of *Af*TFT1-related GT2 glycosyltransferases.

**Table S4**. Glycosidic linkage analysis of oligosaccharide peak fractions collected following HPAEC-PAD separation.

**Table S5**. Summary of lichenase-released oligosaccharides and their molar ratios.

