## Supplementary Figures for "Characterization of a fungal mixed-linkage glucan synthase"

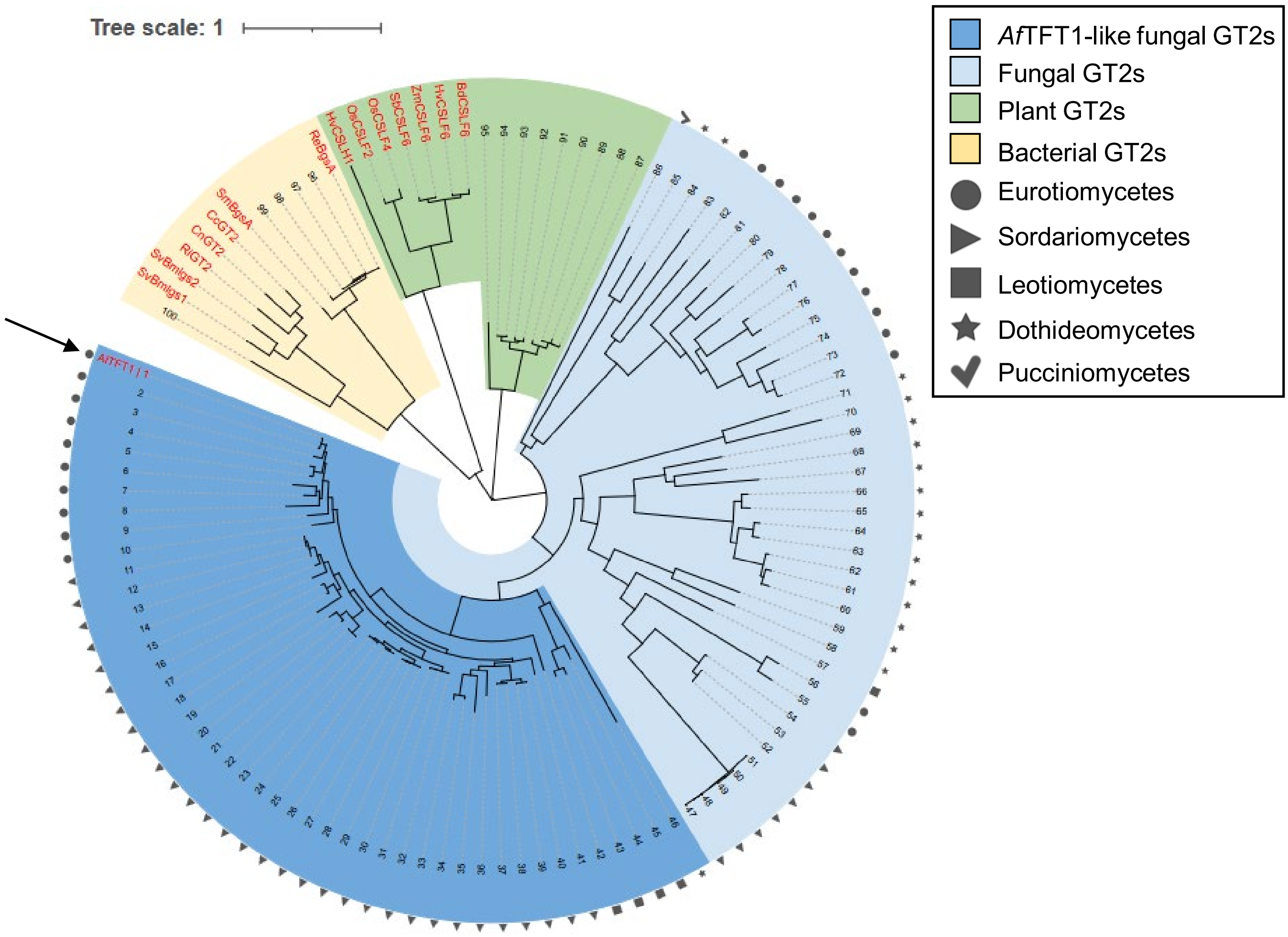


**Supplementary Figure S1: Phylogenetic relationships of AfTFT1-related GT2 glycosyltransferases**
Maximum-likelihood phylogeny of proteins related to *Aspergillus fumigatus* TFT1 together with functionally characterized plant (green), and bacterial (yellow) GT2 proteins. Fungal proteins are indicated in light blue, with *Af*TFT1-like GT2s indicated in dark blue and symbols distinguishing the respective fungal taxonomic groups. Numbers at terminal branches correspond to the BLAST hits listed in Supplementary Table S3. *Af*TFT1 and characterized MLG synthases are highlighted in red; *Af*TFT1 is marked with an arrow. Branch lengths are proportional to the number of substitutions per site (see scale bar).

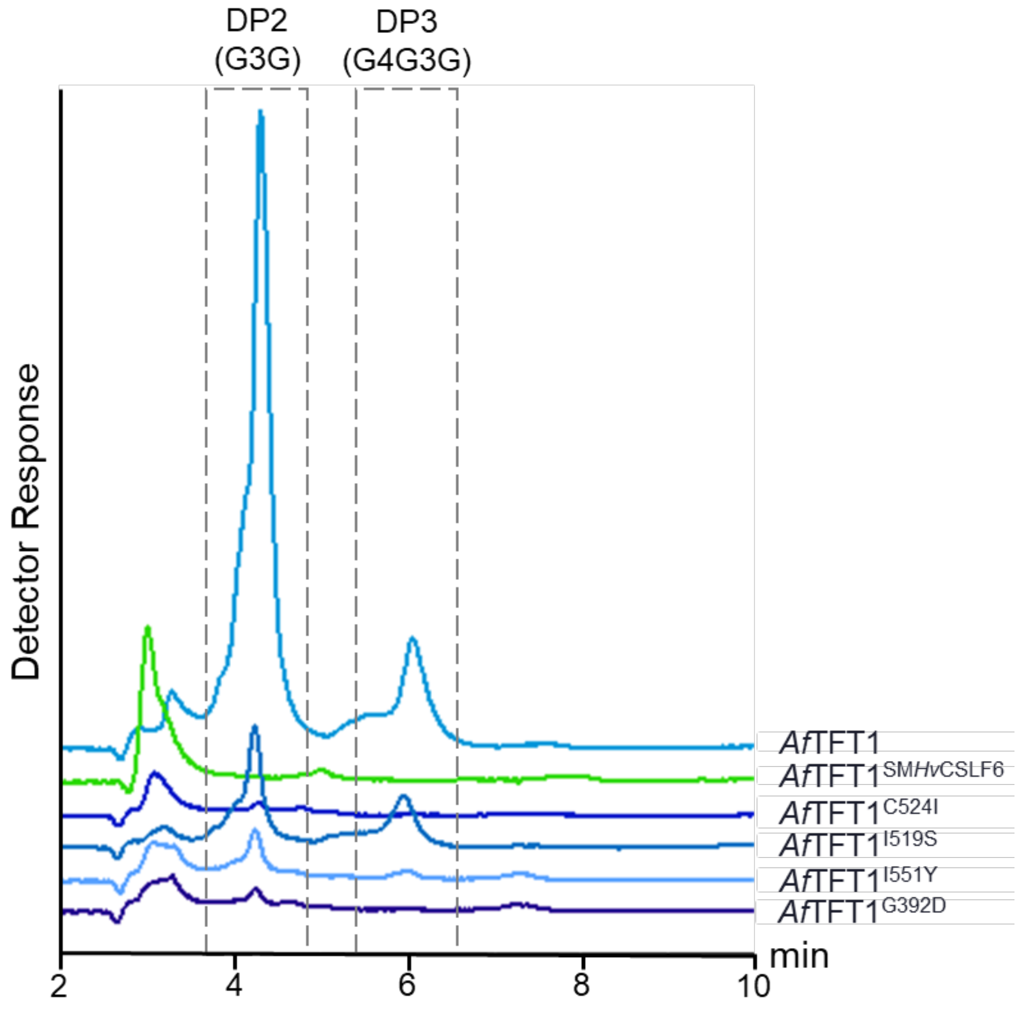


**Supplementary Figure S2: Biochemical characterization of lichenase-released oligosaccharides from *K. phaffii* expressing *Af*TFT1 variants**

Representative HPAEC-PAD chromatograms of lichenase-hydrolysed AIR preparations of *Af*TFT1 cells expressing *Af*TFT1 variants. Released oligosaccharides were identified by comparison with the retention times of MLG-oligosaccharide standards and quantified relative to an internal standard.

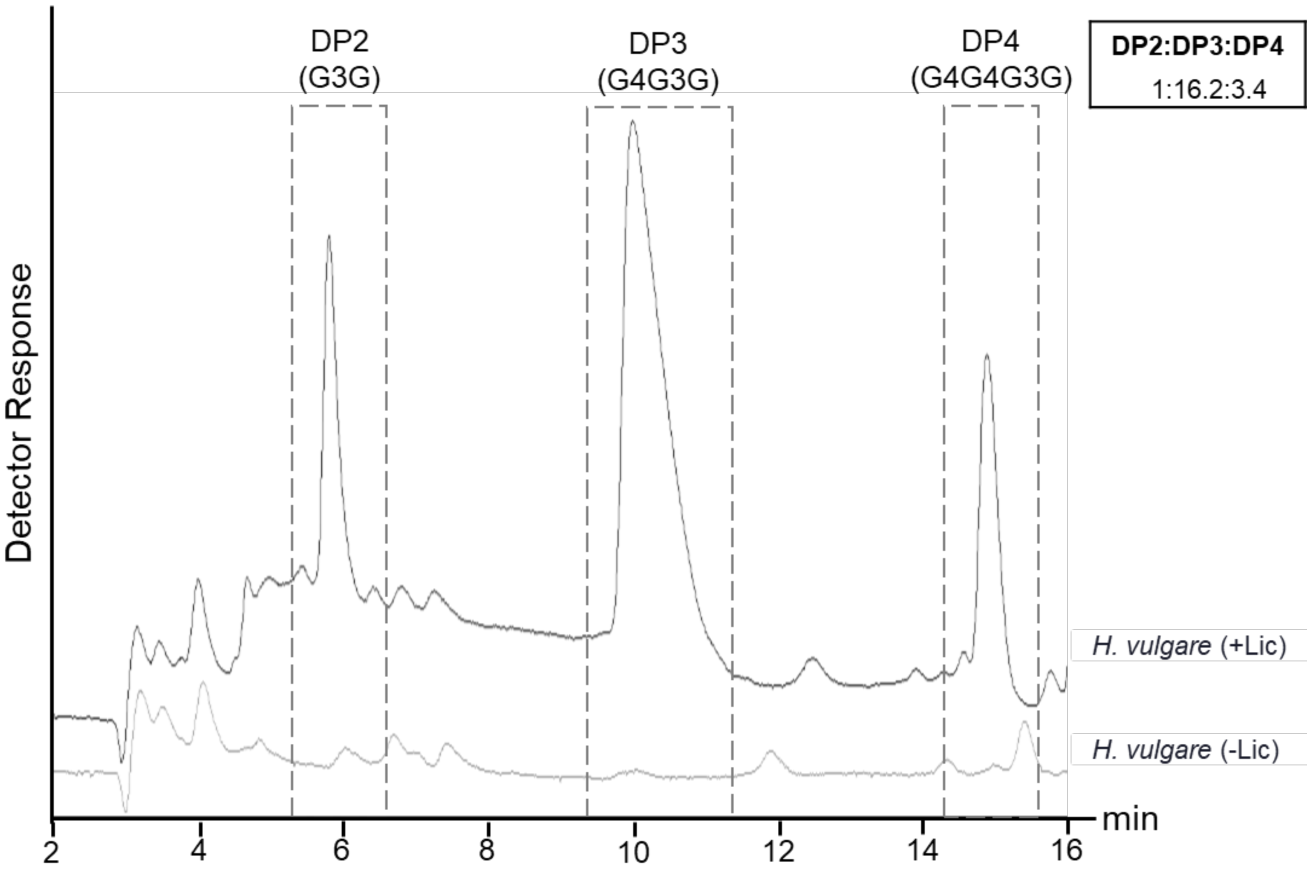


**Supplementary Figure S3: Biochemical characterization of lichenase-released oligosaccharides from *Hordeum vulgare* root material.**

Representative HPAEC-PAD chromatograms of lichenase-hydrolysed AIR preparations of 7-day old *Hordeum vulgare* root material. Released oligosaccharides were identified by comparison with the retention times of MLG-oligosaccharide standards and quantified relative to an internal standard. Inset value indicates the DP2:DP3:DP4 molar ratio of the detected oligosaccharides.
