## Supplementary Tables for "Characterization of a fungal mixed-linkage glucan synthase"

### Supplementary Table S1. Oligonucleotide primers used in this study

| Primer name | Sequence (5´→ 3´) | Purpose | Template/Target |
|---|---|---|---|
| **pAOX1_F** | GCGACTGGTTCCAATTGACAAGC | Verification of integration | pPICZB; *K. phaffii* |
| ***t*AOX1_R** | GGATCCGCACAAACGAAGGT | Verification of integration | pPICZB; *K. phaffii* |
| ***Af*TFT1_F1** | GTAGCACTGATTCGAAATGAACGG AC | Amplification of synthesized *Af*TFT1 | *Af*TFT1 |
| ***Af*TFT1_F2** | GTGCTCGATGACGCCAAGTC | Verification of integration | *Af*TFT1; *K. phaffii* |
| ***Af*TFT1_R1** | ATCGATGCACTCGAGTCAGA | Amplification of synthesized *Af*TFT1 | *Af*TFT1 |
| ***Af*TFT1_R2** | GAGGAAGGACGGCACGCAG | verification of integration | *Af*TFT1; *K. phaffii* |
| ***Af*TFT1$^{SMHvCSLF6}$_F1** | CCTCTCCAGTTCGGTACCGTG | Amplification of synthesized *Af*TFT1$^{SMHvCSLF6}$ | Synthetic fragment *Af*TFT1$^{SMHvCSLF6}$ |
| ***Af*TFT1$^{SMHvCSLF6}$_R1** | CGGATCACGCTCTTCGAGTG | Amplification of synthesized *Af*TFT1$^{SMHvCSLF6}$ | Synthetic fragment *Af*TFT1$^{SMHvCSLF6}$ |
| **Variant_C524I_Fw** | CATTTCCAACCGAATCATCGAGTTC GCCCTCTTC | Generation of Gibson fragment for *Af*TFT1$^{C524I}$ | pPICZB+*Af*TFT1 |
| **Variant_C524I_Rv** | GAAGAGGGCGAACTCGATGATTCG GTTGGAAATG | Generation of Gibson fragment for *Af*TFT1$^{C524I}$ | pPICZB+*Af*TFT1 |
| **Variant_I519S_Fw** | TGCTTTGCCTCGACCTCTTCCAACC GAATCTGCGAG | Generation of Gibson fragment for *Af*TFT1$^{I519S}$ | pPICZB+*Af*TFT1 |
| **Variant_I519S_Rv** | CTCGCAGATTCGGTTGGAAGAGGT CGAGGCAAAGCA | Generation of Gibson fragment for *Af*TFT1$^{I519S}$ | pPICZB+*Af*TFT1 |
| **Variant_I551Y_Fw** | CAGCTCTGGATGGCACCCTACTAT GCC | Generation of Gibson fragment for *Af*TFT1$^{I551Y}$ | pPICZB+*Af*TFT1 |
| **Variant_I551Y_Rv** | GGCA**TA**GTAGGGTGCCATCCAGAG CTG | Generation of Gibson fragment for *Af*TFT1$^{I551Y}$ | pPICZB+*Af*TFT1 |
| **Variant_G392D_Fw** | GGCAACTTCCCTCTTGATTCGTTGG CCGAAGAC | Generation of Gibson fragment for *Af*TFT1$^{G392D}$ | pPICZB+*Af*TFT1 |
| **Variant_G392D_Rv** | GTCTTCGGCCAACGAATCAAGAGG GAAGTTGCC | Generation of Gibson fragment for *Af*TFT1$^{G392D}$ | pPICZB+*Af*TFT1 |
| **Variants_Bb_Fw** | CTGACGCTCAGTGGAACGAAAACT CACGT | Generation of Gibson fragment for *Af*TFT1 variants | pPICZB+*Af*TFT1 |
| **Variants_Bb_Rv** | ACGTGAGTTTTCGTTCCACTGAGC GTCAG | Generation of Gibson fragment for *Af*TFT1 variants | pPICZB+*Af*TFT1 |

**Supplementary Table S2. Protein sequences and accession numbers used for phylogenetic analysis of *Af*TFT1 and characterized MLG synthases**

| | | Polysaccharide | | | | |
|---|---|---|---|---|---|---|
| **Entry** | **Accession** | **Biochemical characterization** | **Evidence** | **Subunits** | **Molar ratio** | **Reference** |
| ***Af*TFT1** | XP_748682.1 | Yes | Structural characterization | DP2; DP3 | 3.4:1 | This work |
| ***Bd*CSLF6** | C5YHD7_SORBI | Yes | Structural characterization | DP3; DP4 | 8:1 | Jobling 2015; Francin-Allami et al. 2023 |
| ***Cc*GT2** | WP_133015619.1 | No | - | - | - | Saldivar et al. 2026 |
| ***Cn*GT2** | WP_066891591.1 | No | - | - | - | Saldivar et al. 2026 |
| ***Hv*CSLF6** | F2DMH9_HORVV | Yes | Structural characterization | DP3; DP4 | 2-3:1 | Taketa et al. 2012; Purushotham et al. 2022 |
| ***Hv*CSLH1** | FJ459581 | Yes | Structural characterization | DP3; DP4 | 3.6:1 | Doblin et al. 2009 |
| ***Os*CSLF2** | CSLF2_ORYSJ | No | Immunolabelling | - | - | Burton et al. 2006 |
| ***Os*CSLF4** | CSLF4_ORYSJ | No | Immunolabelling | - | - | Burton et al. 2006 |
| ***Ri*GT2** | CED93608.1 | Yes | Structural characterization | DP3; DP4 | 2:1 | (Chang et al. 2023) |
| ***Sb*CSLF6** | C5YHD7_SORBI | Yes | Structural characterization | DP3; DP4 | 3.4:1 | Jobling 2015; Kim et al. 2023 |
| ***Sm*BgsA*** | Q92WG2 | Yes | Structural characterization | DP2 | - | Pérez-Mendoza et al. 2015 |
| ***Sv*Bmlgs1** | WP_055257043.1 | Yes | Structural characterization | DP3; DP5; DP7 | - | Lampugnani et al. 2024 |
| ***Sv*Bmlgs2** | WP_055257044.1 | Yes | Structural characterization | DP3; DP5; DP7 | - | Lampugnani et al. 2024 |
| ***Zm*CSLF6** | A0A0R6UPA6_MAIZE | Yes | Structural characterization | DP3; DP4 | 2:1 | Jobling 2015 |

Summary of GT2 domain-containing proteins and MLG synthases included in the phylogenetic tree presented in Figure 2. Details include entry name and database accession number (NCBI/UniProt). "Polysaccharide" column describes characterization of MLG-polysaccharide within its native host organism. Biochemical characterization of the synthase as defined by heterologous expression in a recombinant host system. Asterisk marks that protein has not been biochemically characterized through heterologous expression. Af, *Aspergillus fumigatus*; Bd, *Brachypodium distachyon*; Cc, *Clostridium cuniculi*; Cn, *Clostridium nigeriense*; Hv, *Hordeum vulgare*; Os, *Oryza sativa*; Ri, *Romboutsia ilealis*; Sb, *Sorghum bicolor*; Sm, *Sinorhizobium meliloti*; Sv, *Sarcina ventriculi*; Zm, *Zea mays*.

**Supplementary Table S3. BLAST search results and taxonomic classification of *Af*TFT1-related GT2 glycosyltransferases.**

| Sequence number | *Organism* | Identity | Similarity | Gaps | Protein length | Score (bits) | Description | KEGG gene identifier | KEGG organism code | Major lineage | Taxonomic class |
|---|---|---|---|---|---|---|---|---|---|---|---|
| 1 | *Aspergillus fumigatus* | 100% | 100% | 0% | 727 | 1507 | glycosyl transferase | afm:AFUA_3G03620 | afm | Fungi | Eurotiomycetes |
| 2 | *Aspergillus fischeri* | 96% | 97% | 1% | 738 | 1463 | glycosyl transferase, putative | nfi:NFIA_005820 | nfi | Fungi | Eurotiomycetes |
| 3 | *Aspergillus clavatus* | 85% | 91% | 2% | 739 | 1297 | glycosyl transferase, putative | act:ACLA_060920 | act | Fungi | Eurotiomycetes |
| 4 | *Penicillium digitatum* | 86% | 93% | 0% | 686 | 1200 | Glycosyl transferase, putative | pdp:PDIP_00430 | pdp | Fungi | Eurotiomycetes |
| 5 | *Penicillium rubens* | 79% | 87% | 4% | 753 | 1240 | uncharacterized protein | pcs:N7525_009167 | pcs | Fungi | Eurotiomycetes |
| 6 | *Penicillium psychrofluorescens* | 72% | 83% | 4% | 1109 | 1118 | PFLUO_LOCUS3528; uncharacterized protein | ppsf:300789262 | ppsf | Fungi | Eurotiomycetes |
| 7 | *Penicillium oxalicum* | 84% | 91% | 0% | 775 | 1147 | hypothetical protein | pou:POX_a00964 | pou | Fungi | Eurotiomycetes |
| 8 | *Aspergillus flavus* | 70% | 80% | 3% | 703 | 1048 | hypothetical protein | afv:AFLA_004685 | afv | Fungi | Eurotiomycetes |
| 9 | *Talaromyces rugulosus* | 81% | 89% | 1% | 772 | 1142 | uncharacterized protein | trg:TRUGW13939_01336 | trg | Fungi | Eurotiomycetes |
| 10 | *Talaromyces marneffei* | 71% | 81% | 6% | 800 | 1131 | uncharacterized protein | tmf:EYB26_008787 | tmf | Fungi | Eurotiomycetes |
| 11 | *Metarhizium robertsii* | 74% | 85% | 1% | 777 | 1063 | glycosyltransferase family 2 | maj:MAA_01000 | maj | Fungi | Sordariomycetes |
| 12 | *Metarhizium brunneum* | 74% | 85% | 1% | 777 | 1066 | bcsA, G6M90_00g000620; Cellulose synthase catalytic subunit | mbrn:26244378 | mbrn | Fungi | Sordariomycetes |
| 13 | *Metarhizium acridum* | 74% | 85% | 1% | 777 | 1076 | J3458_021121; uncharacterized protein | maw:19253510 | maw | Fungi | Sordariomycetes |
| 14 | *Pochonia chlamydosporia* | 72% | 83% | 4% | 761 | 1028 | glycosyl transferase | pchm:VFPPC_09949 | pchm | Fungi | Sordariomycetes |
| 15 | *Ustilaginoidea virens* | 77% | 87% | 0% | 786 | 1067 | UV8b_01782; uncharacterized protein | uvi:66062560 | uvi | Fungi | Sordariomycetes |
| 16 | *Purpureocillium takamizusanense* | 73% | 84% | 3% | 769 | 1046 | uncharacterized protein | ptkz:JDV02_004714 | ptkz | Fungi | Sordariomycetes |
| 17 | *Purpureocillium lilacinum* | 74% | 84% | 3% | 768 | 1053 | PLICBS_009827; uncharacterized protein | plj:28893408 | plj | Fungi | Sordariomycetes |
| 18 | *Drechmeria coniospora* | 71% | 84% | 0% | 770 | 979 | DCS_00052; hypothetical protein | dcon:63712695 | dcon | Fungi | Sordariomycetes |
| 19 | *Trichoderma atroviride* | 76% | 87% | 1% | 694 | 964 | TrAtP1_011892; uncharacterized protein | tatv:25783782 | tatv | Fungi | Sordariomycetes |
| 20 | *Trichoderma asperellum* | 69% | 80% | 4% | 755 | 1048 | TrAFT101_008250; uncharacterized protein | tasp:36616334 | tasp | Fungi | Sordariomycetes |
| 21 | *Trichoderma reesei QM6a* | 79% | 88% | 0% | 589 | 989 | hypothetical protein | tre:TRIREDRAFT_77283 | tre | Fungi | Sordariomycetes |
| 22 | *Trichoderma reesei RUT C-30* | 75% | 85% | 0% | 755 | 1046 | hypothetical protein | trr:M419DRAFT_140893 | trr | Fungi | Sordariomycetes |
| 23 | *Fusarium poae* | 79% | 87% | 1% | 821 | 1078 | hypothetical protein | fpoa:FPOAC1_011681 | fpoa | Fungi | Sordariomycetes |
| 24 | *Fusarium venenatum* | 78% | 87% | 1% | 847 | 1084 | uncharacterized protein | fvn:FVRRES_12735 | fvn | Fungi | Sordariomycetes |
| 25 | *Fusarium graminearum* | 78% | 87% | 1% | 824 | 1080 | hypothetical protein | fgr:FGSG_07891 | fgr | Fungi | Sordariomycetes |
| 26 | *Fusarium pseudograminearum* | 77% | 87% | 1% | 851 | 1085 | hypothetical protein | fpu:FPSE_11044 | fpu | Fungi | Sordariomycetes |
| 27 | *Fusarium musae* | 78% | 87% | 1% | 834 | 1080 | hypothetical protein | fmu:J7337_007238 | fmu | Fungi | Sordariomycetes |
| 28 | *Fusarium verticillioides* | 78% | 87% | 1% | 834 | 1083 | hypothetical protein | fvr:FVEG_12066 | fvr | Fungi | Sordariomycetes |
| 29 | *Fusarium oxysporum* | 78% | 87% | 1% | 830 | 1085 | hypothetical protein | fox:FOXG_03792 | fox | Fungi | Sordariomycetes |
| 30 | *Fusarium keratoplasticum* | 80% | 89% | 0% | 847 | 1089 | hypothetical protein | fkr:NCS57_00533600 | fkr | Fungi | Sordariomycetes |

| | | | | | | | | | | | |
|---|---|---|---|---|---|---|---|---|---|---|---|
| 31 | *Fusarium falciforme* | 80% | 89% | 0% | 849 | 1092 | Hypothetical protein | ffc:NCS54_00495800 | ffc | Fungi | Sordariomycetes |
| 32 | *Fusarium vanettenii* | 73% | 83% | 5% | 820 | 1092 | glycosyltransferase family 2 | nhe:NECHADRAFT_73104 | nhe | Fungi | Sordariomycetes |
| 33 | *Akanthomyces muscarius* | 68% | 79% | 4% | 772 | 1027 | hypothetical protein | amus:LMH87_006084 | amus | Fungi | Sordariomycetes |
| 34 | *Cordyceps militaris* | 70% | 80% | 3% | 764 | 1035 | glycosyl transferase, putative | cmt:CCM_05446 | cmt | Fungi | Sordariomycetes |
| 35 | *Sporothrix schenckii* | 71% | 84% | 0% | 843 | 989 | glucosylltransferase family 2 | ssck:SPSK_08488 | ssck | Fungi | Sordariomycetes |
| 36 | *Verticillium alfalfae* | 76% | 84% | 0% | 777 | 1032 | cellulose synthase catalytic subunit | val:VDBG_09908 | val | Fungi | Sordariomycetes |
| 37 | *Colletotrichum destructivum* | 73% | 84% | 2% | 843 | 1061 | CDEST_12666; Putative glycosyltransferase 2, nucleotide-diphospho-sugar transferase | cdet:87949166 | cdet | Fungi | Sordariomycetes |
| 38 | *Colletotrichum higginsianum* | 73% | 84% | 2% | 843 | 1061 | Glycosyltransferase family 2 | chig:CH63R_06180 | chig | Fungi | Sordariomycetes |
| 39 | *Colletotrichum fioriniae* | 73% | 84% | 1% | 843 | 1052 | glycosyltransferase family 2 | cfj:CFIO01_09720 | cfj | Fungi | Sordariomycetes |
| 40 | *Colletotrichum lupini* | 73% | 84% | 1% | 964 | 1054 | glycosyltransferase family 2 | clup:CLUP02_14194 | clup | Fungi | Sordariomycetes |
| 41 | *Phaeoacremonium minimum* | 78% | 87% | 0% | 652 | 1061 | putative glycosyl protein | tmn:UCRPA7_8278 | tmn | Fungi | Sordariomycetes |
| 42 | *Drepanopeziza brunnea* | 68% | 78% | 1% | 874 | 918 | hypothetical protein | mbe:MBM_03149 | mbe | Fungi | Leotiomycetes |
| 43 | *Sclerotinia sclerotiorum* | 51% | 66% | 1% | 789 | 653 | hypothetical protein | ssl:SS1G_10847 | ssl | Fungi | Leotiomycetes |
| 44 | *Botrytis cinerea* | 51% | 67% | 0% | 785 | 665 | hypothetical protein | bfu:BCIN_04g06040 | bfu | Fungi | Leotiomycetes |
| 45 | *Mollisia scopiformis* | 50% | 67% | 0% | 766 | 672 | uncharacterized protein | psco:LY89DRAFT_738802 | psco | Fungi | Leotiomycetes |
| 46 | *Parastagonospora nodorum* | 40% | 60% | 0% | 724 | 491 | hypothetical protein | pno:SNOG_05514 | pno | Fungi | Dothideomycetes |
| 47 | *Podospora pseudocomata* | 30% | 46% | 7% | 725 | 238 | hypothetical protein | ppsd:87908361 | ppsd | Fungi | Sordariomycetes |
| 48 | *Podospora bellae-mahoneyi* | 30% | 46% | 7% | 725 | 237 | hypothetical protein | pbel:QC761_301480 | pbel | Fungi | Sordariomycetes |
| 49 | *Podospora pseudopauciseta* | 31% | 46% | 7% | 725 | 236 | hypothetical protein | ppsp:QC763_301480 | ppsp | Fungi | Sordariomycetes |
| 50 | *Podospora pseudoanserina* | 31% | 46% | 7% | 725 | 237 | hypothetical protein | ppsa:QC764_301480 | ppsa | Fungi | Sordariomycetes |
| 51 | *Podospora anserina* | 30% | 46% | 6% | 557 | 226 | hypothetical protein | pan:PODANSg433 | pan | Fungi | Sordariomycetes |
| 52 | *Podospora pseudopauciseta* | 39% | 57% | 1% | 556 | 288 | hypothetical protein | ppsp:QC763_118300 | ppsp | Fungi | Sordariomycetes |
| 53 | *Podospora pseudocomata* | 39% | 57% | 1% | 556 | 289 | uncharacterized protein | ppsd:87906171 | ppsd | Fungi | Sordariomycetes |
| 54 | *Podospora bellae-mahoneyi* | 35% | 54% | 3% | 633 | 342 | uncharacterized protein | pbel:QC761_118300 | pbel | Fungi | Sordariomycetes |
| 55 | *Talaromyces marneffei* | 34% | 53% | 3% | 644 | 286 | uncharacterized protein | tmf:EYB26_001746 | tmf | Fungi | Eurotiomycetes |
| 56 | *Talaromyces marneffei* | 34% | 53% | 1% | 605 | 305 | uncharacterized protein | tmf:EYB26_007049 | tmf | Fungi | Eurotiomycetes |
| 57 | *Mollisia scopiformis* | 43% | 61% | 1% | 544 | 397 | nucleotide-diphospho-sugar transferase | psco:LY89DRAFT_744124 | psco | Fungi | Leotiomycetes |
| 58 | *Bipolaris sorokiniana* | 38% | 55% | 3% | 543 | 370 | glycosyltransferase family 2 protein | bsc:COCSADRAFT_89237 | bsc | Fungi | Dothideomycetes |
| 59 | *Alternaria alternata* | 42% | 56% | 3% | 572 | 383 | nucleotide-diphospho-sugar transferase | aalt:CC77DRAFT_1082655 | aalt | Fungi | Dothideomycetes |
| 60 | *Bipolaris oryzae* | 38% | 55% | 3% | 620 | 383 | glycosyltransferase family 2 protein | bor:COCMIDRAFT_94307 | bor | Fungi | Dothideomycetes |
| 61 | *Bipolaris zeicola* | 38% | 54% | 5% | 621 | 402 | glycosyltransferase family 2 protein | bze:COCCADRAFT_4124 | bze | Fungi | Dothideomycetes |
| 62 | *Bipolaris sorokiniana* | 37% | 54% | 4% | 621 | 394 | glycosyltransferase family 2 protein | bsc:COCSADRAFT_41996 | bsc | Fungi | Dothideomycetes |

| | | | | | | | | | | | |
|---|---|---|---|---|---|---|---|---|---|---|---|
| 63 | *Alternaria dauci* | 39% | 56% | 5% | 586 | 395 | ACET3X_001314; hypothetical protein | adac:96081636 | adac | Fungi | Dothideomycetes |
| 64 | *Alternaria alternata* | 37% | 53% | 6% | 624 | 405 | cellulose synthase catalytic subunit | aalt:CC77DRAFT_939831 | aalt | Fungi | Dothideomycetes |
| 65 | *Pyrenophora tritici-repentis* | 39% | 55% | 4% | 622 | 405 | PtrM4_082520; glycosyltransferase protein | ptrr:6348413 | ptrr | Fungi | Dothideomycetes |
| 66 | *Pyrenophora teres* | 38% | 54% | 5% | 622 | 405 | hypothetical protein | pte:PTT_07795 | pte | Fungi | Dothideomycetes |
| 67 | *Pseudocercospora fijiensis* | 35% | 52% | 6% | 641 | 377 | glycosyltransferase family 2 protein | pfj:MYCFIDRAFT_157228 | pfj | Fungi | Dothideomycetes |
| 68 | *Baudoinia panamericana* | 39% | 57% | 7% | 603 | 389 | glycosyltransferase family 2 protein | bcom:BAUCODRAFT_138544 | bcom | Fungi | Dothideomycetes |
| 69 | *Baudoinia panamericana* | 36% | 52% | 13% | 649 | 384 | glycosyltransferase family 2 protein | bcom:BAUCODRAFT_29516 | bcom | Fungi | Dothideomycetes |
| 70 | *Cercospora beticola* | 35% | 52% | 3% | 1340 | 244 | Nonribosomal peptide synthetase 13 | cbet:CB0940_12110 | cbet | Fungi | Dothideomycetes |
| 71 | *Cercospora beticola* | 33% | 50% | 5% | 643 | 287 | Cellulose synthase catalytic subunit [UDP-forming] | cbet:CB0940_00365 | cbet | Fungi | Dothideomycetes |
| 72 | *Aspergillus fumigatus* | 31% | 45% | 9% | 621 | 229 | glycosyl transferase | afm:AFUA_8G00680 | afm | Fungi | Eurotiomycetes |
| 73 | *Aspergillus fischeri* | 32% | 46% | 9% | 621 | 231 | glycosyl transferase, putative | nfi:NFIA_094120 | nfi | Fungi | Eurotiomycetes |
| 74 | *Talaromyces marneffei* | 33% | 49% | 7% | 622 | 234 | uncharacterized protein | tmf:EYB26_001943 | tmf | Fungi | Eurotiomycetes |
| 75 | *Aspergillus clavatus* | 31% | 45% | 8% | 556 | 221 | glycosyl transferase, group 2 family protein | act:ACLA_044860 | act | Fungi | Eurotiomycetes |
| 76 | *Aspergillus niger* | 34% | 47% | 8% | 608 | 218 | uncharacterized protein | ang:An03g05740 | ang | Fungi | Eurotiomycetes |
| 77 | *Aspergillus luchuensis* | 33% | 47% | 8% | 614 | 220 | uncharacterized protein | aluc:AKAW2_50941S | aluc | Fungi | Eurotiomycetes |
| 78 | *Aspergillus flavus* | 32% | 44% | 10% | 573 | 208 | hypothetical protein | afv:AFLA_012136 | afv | Fungi | Eurotiomycetes |
| 79 | *Penicillium oxalicum* | 30% | 44% | 5% | 647 | 200 | hypothetical protein | pou:POX_c04080 | pou | Fungi | Eurotiomycetes |
| 80 | *Penicillium psychrofluorescens* | 42% | 55% | 3% | 539 | 199 | PFLUO_LOCUS6476; uncharacterized protein | ppsf:300792536 | ppsf | Fungi | Eurotiomycetes |
| 81 | *Aspergillus nidulans* | 44% | 57% | 3% | 668 | 215 | protein celA | ani:ANIA_08444 | ani | Fungi | Eurotiomycetes |
| 82 | *Aspergillus clavatus* | 31% | 47% | 5% | 623 | 236 | glycosyl transferase, putative | act:ACLA_008150 | act | Fungi | Eurotiomycetes |
| 83 | *Aspergillus niger* | 33% | 51% | 3% | 515 | 196 | uncharacterized protein | ang:An02g05730 | ang | Fungi | Eurotiomycetes |
| 84 | *Cercospora beticola* | 33% | 48% | 4% | 955 | 262 | UDP-glucuronic acid decarboxylase 1 | cbet:CB0940_09209 | cbet | Fungi | Dothideomycetes |
| 85 | *Fulvia fulva* | 34% | 52% | 4% | 636 | 282 | Putative cellulose synthase 3 | ffu:CLAFUR5_03059 | ffu | Fungi | Dothideomycetes |
| 86 | *Melampsora larici-populina* | 28% | 45% | 12% | 954 | 189 | family 2 glycosyltransferase | mlr:MELLADRAFT_118009 | mlr | Fungi | Pucciniomycetes |
| 87 | *Selaginella moellendorffii* | 30% | 47% | 9% | 650 | 209 | GT2A2; glycosyltransferase, CAZy family GT2 | smo:SELMODRAFT_452931 | smo | Plants | Lycopodiopsida |
| 88 | *Selaginella moellendorffii* | 40% | 57% | 4% | 692 | 165 | hypothetical protein | smo:SELMODRAFT_431149 | smo | Plants | Lycopodiopsida |
| 89 | *Selaginella moellendorffii* | 30% | 48% | 10% | 651 | 215 | GT2A6; glycosyltransferase, CAZy family GT2 | smo:SELMODRAFT_452940 | smo | Plants | Lycopodiopsida |
| 90 | *Selaginella moellendorffii* | 30% | 48% | 11% | 662 | 217 | GT2A5/6; glycosyltransferase, CAZy family GT2 | smo:SELMODRAFT_452932 | smo | Plants | Lycopodiopsida |
| 91 | *Selaginella moellendorffii* | 30% | 47% | 9% | 526 | 213 | GT2A3-2; glycosyltransferase, CAZy family GT2 | smo:SELMODRAFT_452928 | smo | Plants | Lycopodiopsida |
| 92 | *Selaginella moellendorffii* | 30% | 46% | 6% | 661 | 217 | GT2A3; glycosyltransferase, CAZy family GT2 | smo:SELMODRAFT_452938 | smo | Plants | Lycopodiopsida |
| 93 | *Selaginella moellendorffii* | 31% | 47% | 9% | 659 | 219 | GT2A1-2; glycosyltransferase, CAZy family GT2 | smo:SELMODRAFT_452933 | smo | Plants | Lycopodiopsida |
| 94 | *Selaginella moellendorffii* | 31% | 48% | 10% | 659 | 220 | GT2A1; glycosyltransferase, CAZy family GT2 | smo:SELMODRAFT_452930 | smo | Plants | Lycopodiopsida |

| | | | | | | | | | | | |
|---|---|---|---|---|---|---|---|---|---|---|---|
| 95 | *Physcomitrium patens* | 29% | 46% | 10% | 671 | 186 | uncharacterized protein isoform X1 | ppp:112289633 | ppp | Plants | Bryopsida |
| 96 | *Rhizobium sp. N731* | 32% | 47% | 10% | 664 | 165 | cellulose synthase subunit CelA protein | rhx:AMK02_PE00382 | rhx | Bacteria | Alphaproteobacteria |
| 97 | *Rhizobium esperanzae* | 32% | 47% | 10% | 664 | 165 | cellulose synthase subunit CelA protein | rez:AMJ99_PC00382 | rez | Bacteria | Alphaproteobacteria |
| 98 | *Rhizobium etli bv. mimosae Mim1* | 33% | 47% | 10% | 664 | 164 | cellulose synthase CelA-like protein | rel:REMIM1_PD00382 | rel | Bacteria | Alphaproteobacteria |
| 99 | *Rhizobium sp. SRDI969* | 33% | 47% | 10% | 664 | 166 | glycosyltransferase | rhis:N2A41_29530 | rhis | Bacteria | Alphaproteobacteria |
| 100 | *Floricoccus penangensis* | 29% | 46% | 11% | 670 | 162 | glycosyltransferase | fpen:KIW23_00420 | fpen | Bacteria | Bacilli |

| Entry | Organism | Identifier | Major lineage |
|---|---|---|---|
| ***Bd*CSLF6** | ***Brachypodium distachyon*** | **I1GL52_BRADI** | **Plants** |
| ***Cc*GT2** | ***Clostridium cuniculi*** | **WP_133015619.1** | **Bacteria** |
| ***Cn*GT2** | ***Clostridium nigeriense*** | **WP_066891591.1** | **Bacteria** |
| ***Hv*CSLF6** | ***Hordeum vulgare*** | **F2DMH9_HORVV** | **Plants** |
| ***Hv*CSLH1** | ***Hordeum vulgare*** | **FJ459581** | **Plants** |
| ***Os*CSLF2** | ***Oryza sativa*** | **CSLF2_ORYSJ** | **Plants** |
| ***Os*CSLF4** | ***Oryza sativa*** | **CSLF4_ORYSJ** | **Plants** |
| ***Re*BgsA** | ***Rhizobium etli*** | **Q2K103** | **Bacteria** |
| ***Ri*GT2** | ***Romboutsia ilealis*** | **CED93608.1** | **Bacteria** |
| ***Sb*CSLF6** | ***Sorghum bicolor*** | **C5YHD7_SORBI** | **Plants** |
| ***Sm*BgsA** | ***Sinorhizobium meliloti*** | **Q92WG2** | **Bacteria** |
| ***Sv*Bmlgs1** | ***Sarcina ventriculi*** | **WP_055257043.1** | **Bacteria** |
| ***Sv*Bmlgs2** | ***Sarcina ventriculi*** | **WP_055257044.1** | **Bacteria** |
| ***Zm*CSLF6** | ***Zea mays*** | **A0A0R6UPA6_MAIZE** | **Plants** |

Summary of protein sequence similarity searches, including sequence identifiers, source organisms, sequence identity and similarity, alignment gaps, protein length, bit score, and sequence descriptions. KEGG gene identifiers and organism codes, together with major lineage and taxonomic class, are provided for taxonomic classification of all analysed proteins. Additional reference protein entries—including functionally characterized GT2 enzymes added manually for comparative phylogenetic analysis—are also included. Rows are color-coded according to the taxonomic origin of the entries: plants (green), bacteria (yellow), fungi (blue), and oomycetes (purple).

**Supplementary Table S4. Glycosyl linkage analysis of oligosaccharide peak fractions collected following HPAEC-PAD separation.**

| | Collected fractions | | MLG Standards | |
|---|---|---|---|---|
| | **DP2** | **DP3** | **DP2 (G3G)** | **DP3 (G4G3G)** |
| **t-Glc** | **41.0 ± 2.4** | **24.0 ± 2.5** | **42.0** | **26.7** |
| **3-Glc*p*** | **59.0 ± 2.4** | **28.0 ± 2.8** | **58.0** | **35.0** |
| **4-Glc*p*** | | **48.0 ± 3.7** | | **38.3** |

Oligosaccharide peak fractions generated by lichenase hydrolysis of recombinant *K. phaffii* AIR were collected individually following HPAEC-PAD separation and subjected to glycosyl linkage analysis. Data are presented as relative molar percentage (mean ± SD) calculated from three technical replicates. Authentic oligosaccharide standards were analysed in parallel as controls.

**Supplementary Table S5. Summary of lichenase-released oligosaccharides and their molar ratios.**

| | DP2 (G3G) [ng/mg AIR] | DP3 (G4G3G) [ng/mg AIR] | DP4 (G4G4G3G) [ng/mg AIR] | released MLG [ng/mg AIR] | Molar ratio |
|---|---|---|---|---|---|
| ***A. fumigatus*** | 110 ± 9.6 | 48.9 ± 7.4 | n.d. | 159.2 ± 16.7 | 3.4:1 |
| ***Af*TFT1** | 12100.9 ± 3234.6 | 4160.5 ± 1071.7 | n.d. | 16261.5 ± 4019.8 | 2.9:1 |
| ***Hordeum vulgare*** | 75.7 ± 31.7 | 1789.6 ± 222.7 | 491.4 ± 61.9 | 2356.7 ± 282.9 | 1:16.2:3.4 |
| ***Af*TFT1$^{SMHvCSLF6}$** | n.d. | n.d. | n.d. | n.d. | n.a. |
| ***Af*TFT1$^{C524I}$** | 7.7 ± 7.7 | 11.7 ± 15.3 | n.d. | 19.4 ± 21.4 | 1:1 |
| ***Af*TFT1$^{I519S}$** | 13.3 ± 3.7 | 19.4 ± 21.4 | n.d. | 33.3 ± 9.2 | 1:1 |
| ***Af*TFT1$^{I551Y}$** | 1.7 ± 0.3 | n.d. | n.d. | 1.7 ± 0.3 | n.a. |
| ***Af*TFT1$^{G392D}$** | 227.9 ± 9.6 | 392.1 ± 214.4 | n.d. | 620.1 ± 303.4 | 1:1.2 |

Lichenase-released oligosaccharides detected across all samples analysed in this study are summarized together with their corresponding molar ratios. DP, degree of polymerization; n.a., not applicable; n.d., not detected.
